# AFM Microfluidic Cantilevers as Weight Sensors for Live Single Cell Mass Measurements

**DOI:** 10.1101/2022.02.21.481347

**Authors:** Chen-Chi Chien, Jiaxin Jiang, Bin Gong, Tao Li, Angelo Gaitas

## Abstract

Reliably measuring small mass changes at the single-cell level is challenging. In this manuscript, we report the use of microfluidic cantilevers in liquid with sub-nanogram scale weight sensing capability for the measurement of cellular mass changes of living single cells. With this instrumentation, we were able to perform fast mass measurements within 3 minutes. We show results of mass measurements of polystyrene and metal beads of various sizes (smallest weight measured at 280 ± 95 pg) and live single-cell mass measurements in a physiologically relevant environment. We also performed finite element analysis to simulate and optimize the structural design and materials of cantilevers. Our simulation results indicate that using polymer materials, such as SU8 and polyimide, could improve the minimal detectable mass by 3-fold compared to conventional silicon cantilevers. The simulations also suggest that smaller dimensions of length, width, and thickness would improve the mass detection capability of microfluidic cantilevers.

## 1. Introduction

Changes in cell mass provide insights into cellular processes, such as growth patterns, metabolism, migration, and proliferation(1–3), and are indicative of cellular health (4–12). Most existing tools rely on volumetric measurements of cells, as the volume is an indirect measure of biomass, with inaccuracies arising from osmotic changes and other processes in single cells(3, 13, 14). Cell masses range from several nanograms to a few picograms, for example, a yeast cell mass was measured at 11 pg(15). In addition, mass changes due to cellular processes can occur within milliseconds (16) making both mass and temporal resolution essential.

The resonance frequency change of a microcantilever can be used to measure changes in mass (17–20). The resonance frequency, *f_o_*, is a function of the spring constant, which depends on the Young’s modulus and cantilever dimensions, and the effective mass of a resonating microcantilever. Therefore, a cell mass added to a microcantilever can be measured based on changes to the resonance frequency(21, 22). This principle has been used to detect single-cell mass using microchannel resonators(23), pedestal sensors(24), and conventional functionalized atomic force microscopy microcantilevers(3, 25).

Microfluidic atomic force microscopy (AFM) cantilevers, or microfluidic cantilevers in short, which have an embedded fluidic channel along its length and an aperture at the tip, were initially used for fountain-pen lithography applications(26–29). Microfluidic cantilevers were first used for single-cell applications by Meister et al. (30). Since then, these devices have been used for a number of biologically relevant applications(30–36), such as liquid delivery and adhesion measurements. Furthermore, microfluidic cantilever devices with and without tips of various materials have been reported and used for a wide range of applications such as SU8 and 3D printed cantilevers (26,30,37–46).

There are several advantages to use microfluidic cantilevers for mass measurement applications compared to functionalized AFM microcantilevers(3, 25). First, cells can be attached to the tip by applying a negative pressure, which enables the study of both adherent and non-adherent cells. This feature also allows for the cantilevers to be reused immediately after a measurement since measured cells can be easily removed and new cells attached by modulating the pressure, thus enabling the measurement of multiple cells within a short time(47). Finally, the location of cell attachment is always fixed at the channel opening of the microfluidic cantilevers, eliminating the tedious work of determining the locations relative to the tip, which adds complexity to mass calculation^48^. Preliminary results demonstrated the use of microfluidic cantilevers to measure the mass of yeasts and beads in air down to tens of picograms(48).

In this work, we are reporting, for the first time, measurements of the mass of single live cells in a liquid environment with microfluidic cantilevers. Various aspects of this mass measurement scheme are discussed. We measured microspheres with known mass to calibrate the microcantilevers with the thermal tune method (also referred to as the thermal noise, or Sader method) (49, 50), with smallest polystyrene beads of 8 μm measured to be 280 ± 95 pg. We extend the applications to single cell measurements and discuss the limitations of the instrumentations. In addition, finite element analysis (FEA) is used to study resonating microcantilevers in liquid environments in order to find optimal structural material selection and geometry for the purpose of achieving an even lower minimum detectable mass as a future research direction.

## 2. Results and Discussion

### Experiments and Methods

The experimental set-up shown in ***Fig. 1a*** includes a microfluidic cantilever (Cytosurg AG, Switzerland) with an aperture at the tip submerged under liquid, where targets of interest, such as microspheres and living cells, reside. The thermal resonance frequency was measured by Sader method using a Flex-Bio AFM (Nanosurf AG, Switzerland). The AFM is equipped with an inverted microscope, Axio Observer (Carl Zeiss, Germany), and an environmental control enclosure. A pump (Cytosurge AG, Switzerland) is connected to the cantilever and is able to apply pressure from −800mbar to 1000mbar.

**Fig. 1.**
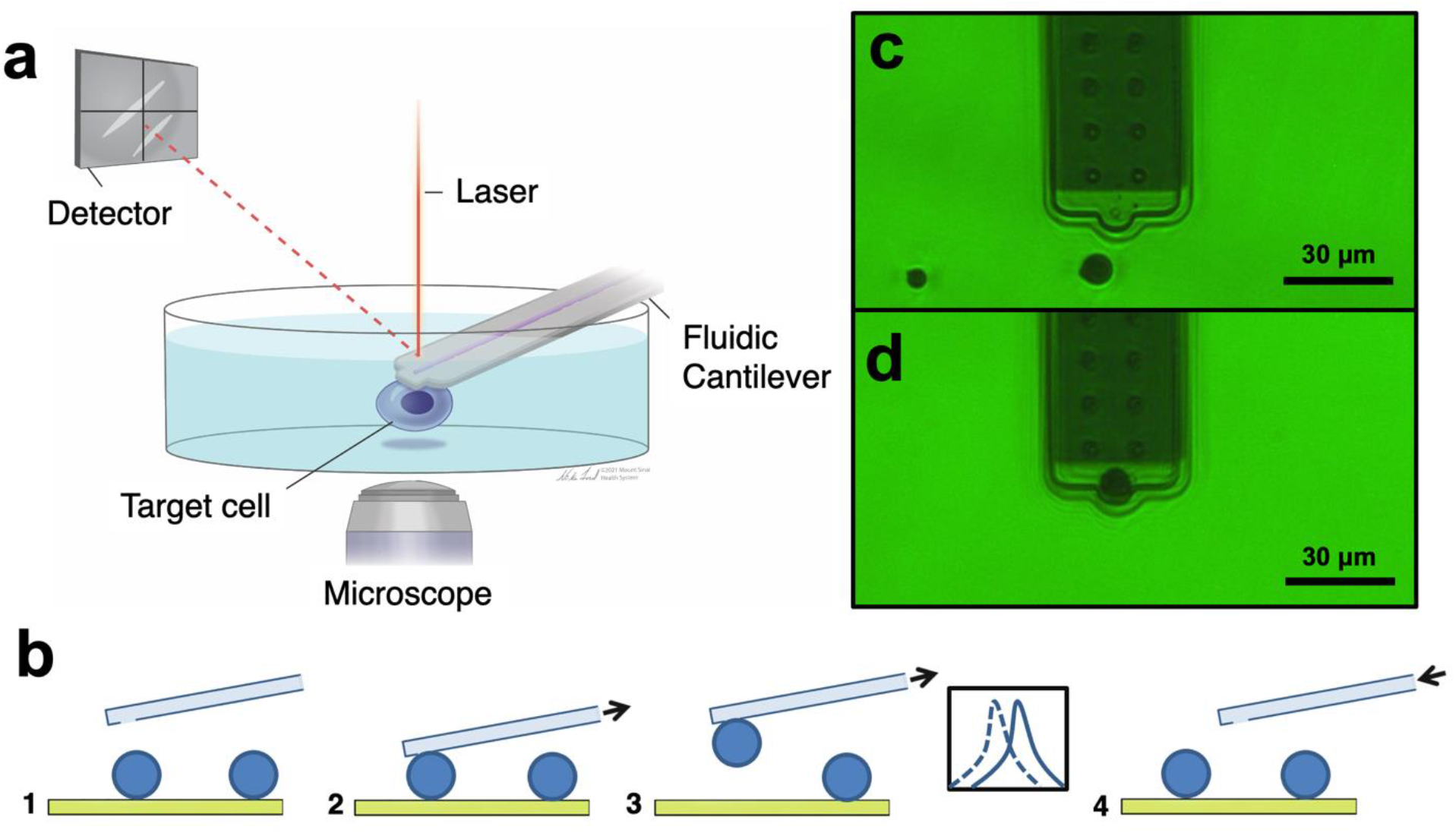
a) Schematic of a microfluidic cantilever for mass measurement set-up. b) Schematics of the flow of mass measurement. (1) First, we identify the target of interest, and move the cantilever close to the target. (2) When the target is sufficiently close to the cantilever, a negative pressure is applied to create a suction to attach the target. (3) The target, now attached to the cantilever, is being raised as far as possible from the surface while still in liquid, and resonance frequency measurement could be performed. (4) After the measurement, a positive pressure is applied to eject the target. The resonance frequency is measured again for subsequent calculation. The cantilever then returns to its initial position and moved to measure another target. c) Microfluidic cantilever approaching the target (metal microsphere), d) The target (metal microsphere) is picked up by the microfluidic cantilever.

The flow of the measurement is illustrated in ***Fig. 1b***. First, the cantilever is moved from its initial home position to the target, and negative pressure is applied to create a suction force to attach the target. The resonance frequency of the cantilever is recorded both with and without the target. Mass can be calculated from change in frequency peak using the equation(21, 22):

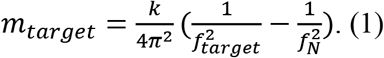

m_target_ is the measured mass of the target, k is the spring constant of the cantilever, and f_N_ and f_target_ are the resonance frequencies in liquid without and with target attachment, respectively. Positive pressure is applied to detach the target, allowing for the immediate measurement of another target. If the target is not ejected by the positive pressure, the cantilever can be gently lifted out of the liquid so that the target is removed by the surface tension of the liquid. This process allows us to reuse the same microfluidic cantilever for multiple measurements following Cytosurge protocols. In general, the device can be reused for 3-5 experiments and up to 3 weeks. Pressure controlled capture mechanism eliminates the needs for adhesion coating of the cantilevers. A microfluidic cantilever with a 4 μm aperture submerged under DI water is shown in ***Fig. 1c***. A metal microsphere was picked up at the tip of the microfluidic cantilever, from the petri dish by applying a pressure of −800mbar as shown in ***Fig. 1d***. We were able to attach, measure, and remove targets within 3 minutes and then move on to the next target.

The thermal tuning method is used to determine the resonance frequency (f_0_) of the cantilever and the quality factor of the cantilever - Q - defined as *fo*/Δ*f* where Δ*f* is the resonance peak width at half maximum as shown in ***Fig. 2b***. The measured resonance frequency, determined by fitting the spectral density to the Lorentzian curve, as shown in ***Fig. 2a***, is measured in air to determine the spring constant of the cantilever while having higher Q factor (Q~100). When the cantilever is submerged in the liquid, the Q factor is decreased (Q~4) due to damping from liquid, yet we could still identify the peak of the resonance frequency as shown in the blue curve in ***Fig. 2c*** and ***Fig. 2d*** by averaging over multiple readout iterations. The decrease of the resonance frequency when an object is attached is shown in the red curves in ***Fig. 2c*** and ***Fig. 2d*** The spring constant was measured in air, and used for later calculations to determine the mass of the target using **Eq. 1** (22).

**Fig. 2.**
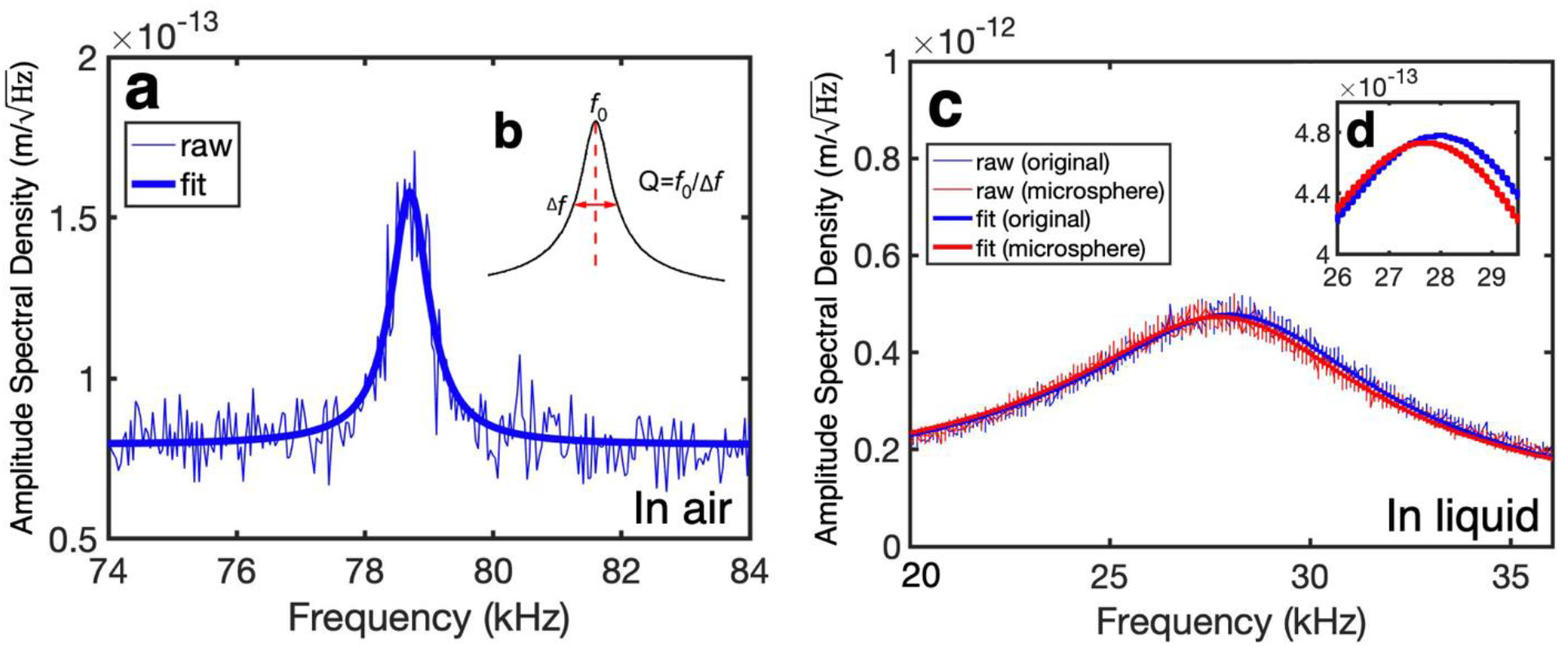
Amplitude spectral density versus frequency measured a) in air, and c) in liquid. The blue curves are the initial spectral density curve and its fit curve without any target attached, and the red curve shows a spectral density curve and its fit curve with a microsphere attached to the cantilever as in ***Fig. 1d***. b) The inset schematic demonstrates the relationship between f_0_, Δf, and Q. d) The enlarged fit curves in liquid, and the change in peak is clearly identified.

We conducted measurements within a petri-dish filled with DI water or growth media using a tipless microfluidic cantilever with openings ranging from 2 μm to 8 μm, and the spring constant of the cantilever ranging between 1.7 N/m and 2.62 N/m. The pump applied <20 mbar positive pressure until the cantilever approached the target. Then negative pressure of −300 to −800 mbar was applied for attachment. A 1,000 mbar of positive pressure was applied for removal after each experiment. To understand the accuracy and limitations of the microfluidic cantilever, we measure the mass of size-standard polystyrene microspheres diameters of 8±0.09,10±0.09, 12±0.1, and 15±0.12μm (NIST size standard, Thermo Scientific 4000, USA), with the diameters and errors provided by the manufacturers. For each diameter of spheres, 4 to 6 spheres are measured. The results are shown in ***Fig. 3a***. The smallest frequency shift measured for a particular 8 μm microsphere was 42Hz, corresponding to the measured mass of approximately 280pg. The expected mass is calculated by assuming a uniform sphere with mass:

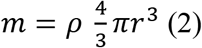

where *ρ* the density of polystyrene, is 1.05 g/cm^3^, and r is the radius of the sphere. The dotted line represents the measured mass in perfect agreement with the estimated mass. The measured mass is within 100 pg of the estimated mass, and the standard deviation of the measurement is within 100 pg, showing a good agreement between the mass measured and the theoretical estimated mass. Despite the fact that **Eq. 1** is an approximation, and is not entirely accurate in liquid,(22) the results we obtained highlight that targets weighing as low as two hundred picograms could be measured. Other comprehensive analytical models for the mass measurements in liquid can further improve the accuracy of measurements of mass in liquid.(51) Other potential errors are attributed to the estimation of spring constant, the fitting of the resonance frequency curves due to thermal noise, a lower Q factor in liquid (***Supplementary Information 1,2***), and the variations of sphere sizes by the manufacturer. Nonetheless, the results allow us to establish a picogram weight sensing system that could be used for measuring small masses.

**Fig. 3.**
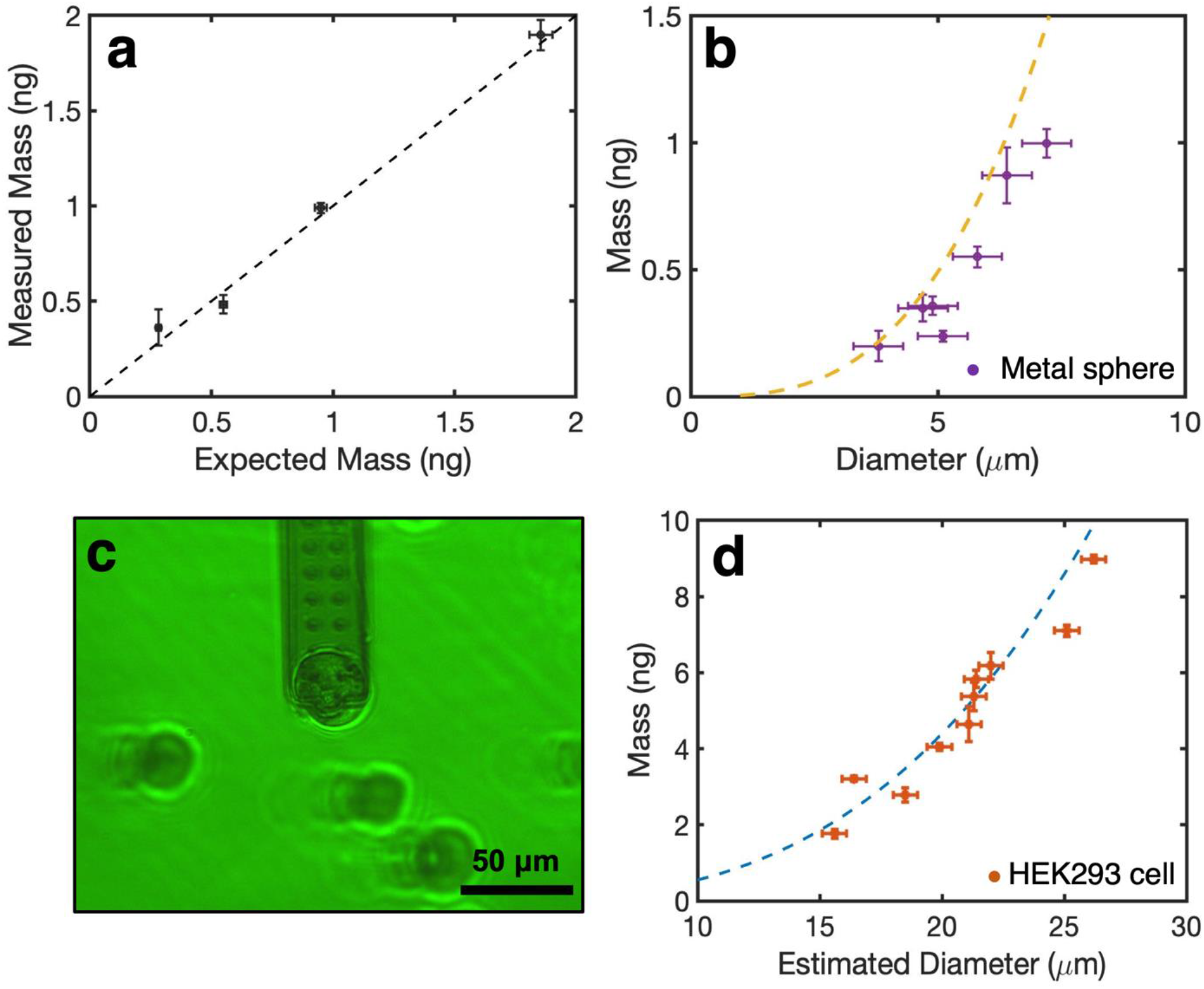
a) Measurements of the masses of size-standard polystyrene microspheres with manufacturer-specified diameters of 8,10,12,15um, compared with their expected masses. The dotted line corresponds to where the measured mass perfectly matches the estimated mass. b) Masses of 7 stainless-steel metal microspheres with diameters from 3.8 to 7.2 μm. The yellow dotted curve is the theoretical mass based on the measured diameters. The error bar for the measured estimated diameter is ± 1 μm. c) A HEK293 cell is picked up by the microfluidic cantilever. d) Summary of masses of 10 HEK293 cells in media with diameter between 15.6 to 26.2 um. The dotted blue curve is the estimated mass of the cells based on the measured diameters. The error bar for the measured estimated diameter is ± 1 μm.

Masses of stainless-steel metal microspheres (Cospheric, USA) with density of 7.8 g/cm^3^ and sphere diameters measured to be from 3.8 to 7.2 μm were also measured and shown in ***Fig. 3b***. Each sphere was measured up to five times. The theoretical mass based on the measured diameter for the microspheres is shown in the yellow dotted curve. We measured the masses of the metal microspheres to be 0.2–1.0 ng for diameters ranging between 3.8 and 7.2 μm. The masses measured largely fall along the theoretical predicted curve, demonstrating, again, that the microfluidic cantilever is measuring the masses of the metal spheres accurately. In addition to the errors cited above, the deviation from the predicted curve can also result from errors of the optical determination of size. The size of the metal spheres was determined using an optical picture and computer software (Gwyddion, Czech Republic). The error for the measured estimated diameter is ± 1 μm as shown in the error bar in ***Fig. 3b***. A more accurate measurement and characterization of the spheres in 3D could reduce the error.

The microfluidic cantilever was also used to measure single live cells. We measured the mass of HEK293 cells in HEPES buffered growth media. (***Supplementary Information 3***) The measuring environment was maintained at 37 °C. Similar to above, the pump was kept at 20mbar positive pressure until the target HEK293 cell was in close proximity to the microfluidic cantilever. Then a negative pressure of −300 mbar was applied to attach the cell. After attachment, the pressure was reduced to −100bar to minimize possible cell disturbance, and the resonance frequency was measured. The different pressures applied do not affect the measured resonance frequency as shown in the ***Supplementary Information 4***. After each measurement, a 1000 mbar of positive pressure was applied for cell removal. Often cells are difficult to remove via pressure alone. In these cases, the cantilever was gently lifted out of the media. The liquid surface tension was strong enough to detach the cells. Following cell removal, the experiment was repeated with another cell. An attached HEK293 cell on the cantilever is showed ***Fig. 3c***. We measured 10 HEK293 cells in media with masses between 1.8 to 9.0 ng and diameters between 15.6 μm to 26.2 μm. The cell mass values measured are consonant with those obtained with using other methods(52). We estimated the volume and correspondingly the mass of the cells assuming the cells are spheres with density of 1.05 gm/cm^3^ using **Eq. 2**. The results are summarized in ***Fig. 3d***. The dotted blue curve represents the estimated mass of the cells based on the measured diameters. It can be observed that the experimental values are very close to the estimated values. In addition to the possible errors outlined previously, another source of the discrepancy could stem from the fact that the cells are not perfectly spherical and therefore our approximation of diameter is not entirely accurate. The error bar for the measured estimated diameter is ± 1 μm as shown in ***Fig. 3d.***

When having the cells attached for 10 minutes, we observed that the mass of some cells remained relatively the same, while for others there were a significant mass loss. ***Fig 4a*** shows the mass variations of several cells attached to the cantilever measured in 5-minute intervals. Two cell masses (* and x) remained relative stable, while (+ and o) had a decrease in mass. A polystyrene microsphere (•) of 12 *μ*m was used as a control. ***Fig. 4b*** shows the mass decrease of cell (o) and its corresponding images at 0, 5, and 10 minutes. We observed optically that the cell diameter shrunk as shown in ***Fig. 4c, d, e***. We used the measured mass and diameter with **Eq. 2** to estimate the density of the cell at different times, and we observe the density of the cell increased from 0.86 g/cm^3^ to 1.17 g/cm^3^ within a 10-minute period. We carefully controlled and performed isolated experiments to rule out suction from the cantilever being the cause. Changes in osmotic pressure (salinity) due to evaporation of media might causes the loss in weight of these cells but further investigation is required to confirm this hypothesis and other causes for the weight reduction. (***Supplementary Information 5&6***). A temperature and moisture controlled enclosed chamber could be utilized to alleviate the issue of media evaporation and improve measurement consistency.

**Fig. 4.**
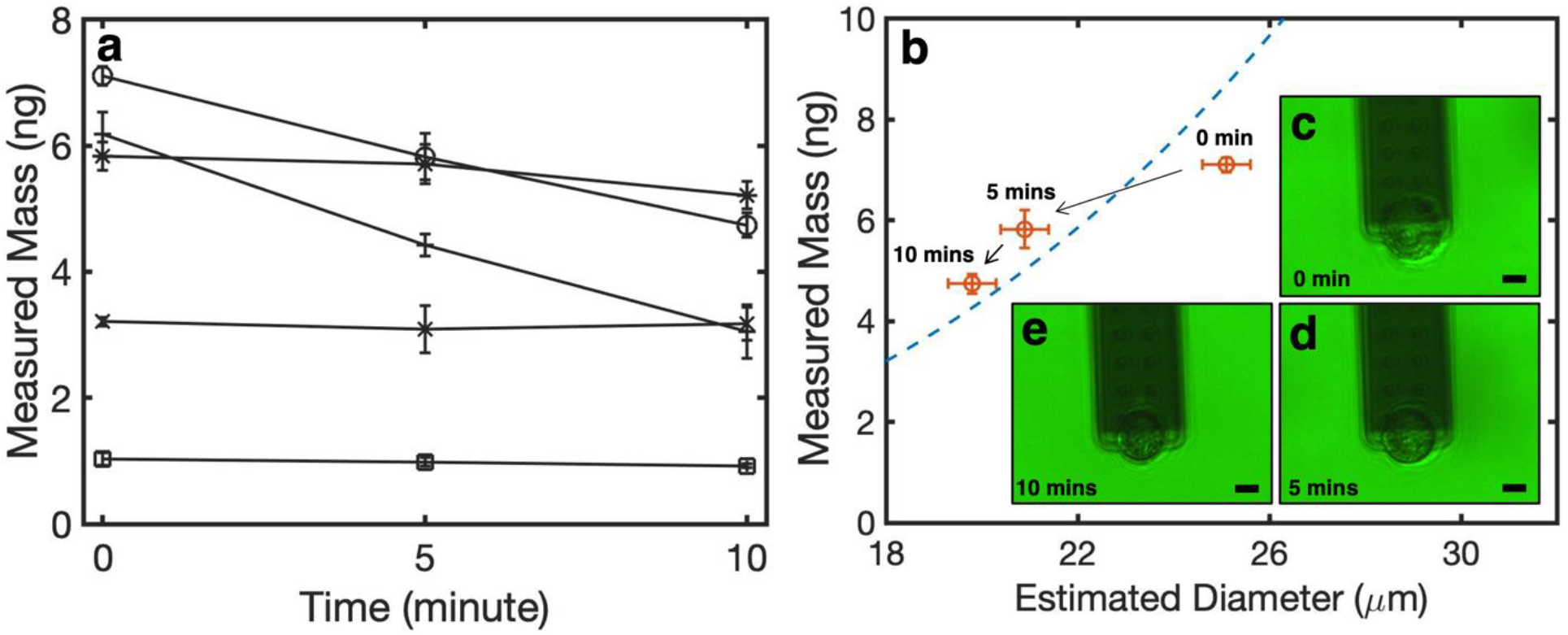
a) Four different HEK293 cells (o, *, +, and x), and a polystyrene microsphere (•) and their masses being measured across 10 minutes time frame. b) The measured mass and diameter for cell (o) at 0, 5, and 10 minutes. c) to e) The optical images of the cell (o) at 0, 5, 10 minutes respectively. The scale bars indicate a length of 10 *μ*m.

### Finite Element Analysis

We conducted finite element analysis (FEA) in order to further optimize the mass sensing capability of the microfluidic cantilever, and to understand its limitations. The resonant characteristics of the cantilever, specifically f_0_ and Q, are mainly determined by the structural and material properties of the cantilever as well as the damping effect. A resonator with a higher quality factor is highly desirable to provide a sharper peak at the resonant frequency, which can lead to a better resolution for signal readout and thus higher measurement accuracy.

While the quality factor in vacuum is primarily limited by internal structural damping of the cantilever such as thermoelastic damping, the resonance of a cantilever in a medium, such as air or water, is heavily affected by viscous damping of the medium (53). Finite element analysis using COMSOL Multiphysics® was conducted to evaluate the resonant characteristics of the cantilever (length 450 μm, width 50 μm, thickness 2 μm) in air and water. The FEA model and the resulting resonant characteristics are shown in ***Fig. 5***.The simulation was based on the thermoviscous acoustics (frequency domain) simulation module in COMSOL Multiphysics, which was used to model both thermal and viscous damping effects of the resonating cantilever in the medium. The parameters used in the simulation are listed in ***Supplementary Information 7***. As shown in ***Fig. 5b***, the simulated f_0_ and Q both match well with experimental results performed with the Si cantilever of the same dimensions (Stat0.2LAuD-10, Nanosurf, Switzerland), and we see that the Q is much lower in the liquid compared to in air.

**Fig. 5.**
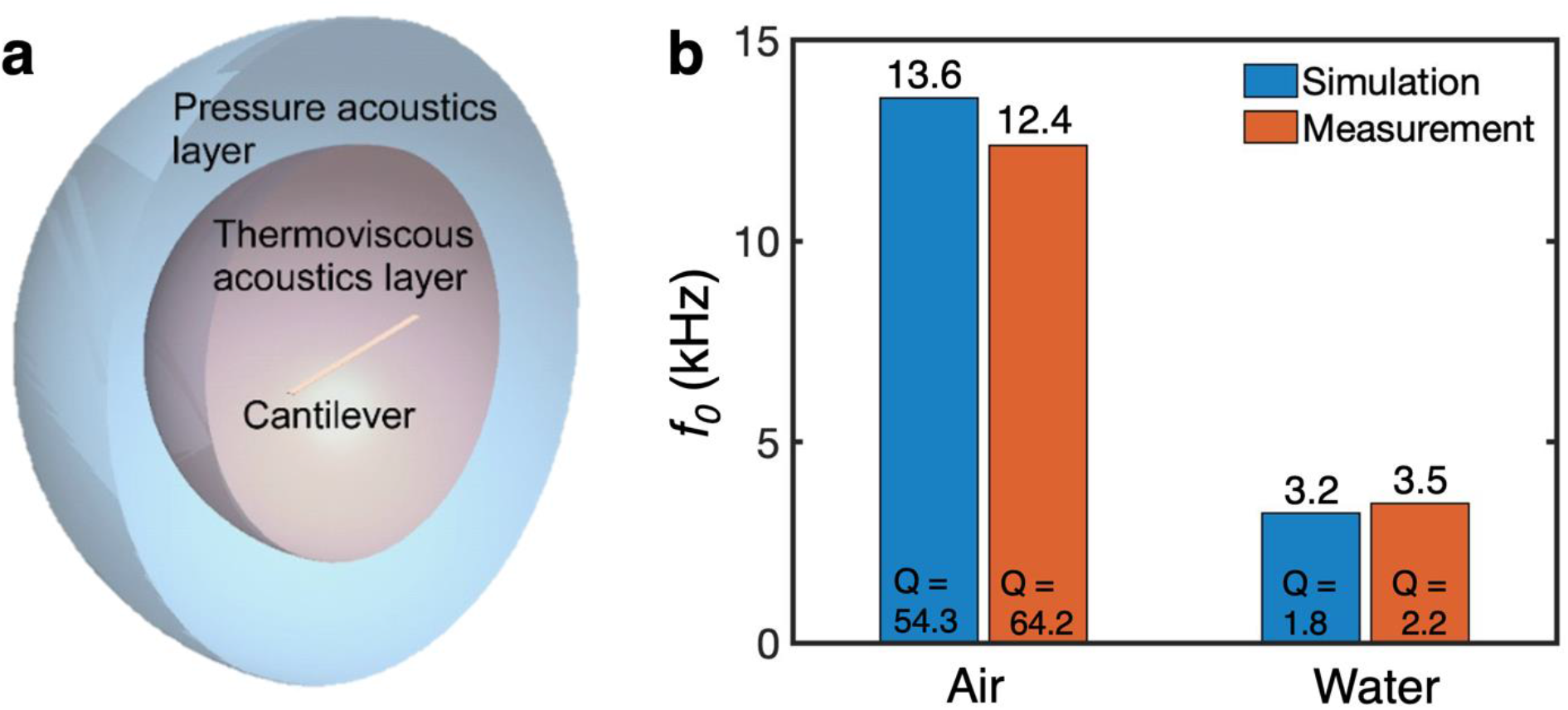
a) Simulation model; b) comparison of resonant characteristics from simulation and experiments for a Si cantilever.

The resonant frequency *f_0_* in liquid can be calculated with **Eq. 3**(54):

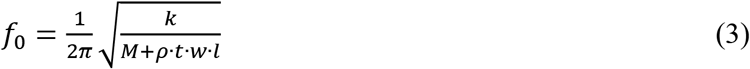

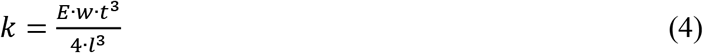

where *E* is the Young’s modulus of the cantilever in Pa, *M* is the mass added to the free end of the cantilever in kg, *ρ* is the density of the cantilever material in kg/m^3^, and *t, w*, and *l* are the thickness, width, and length of the cantilever in m, respectively. When the cantilever dimensions and surrounding medium are kept unchanged while considering different cantilever material options, *f_o_* becomes proportional to 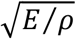.

The minimum detectable mass (i.e., mass resolution) can be characterized by minimum detectable frequency shift δ(Δ*f*) and the cantilever mass sensitivity *S_m_*. The minimum detectable frequency shift δ(Δ*f*) is limited by the thermal noise and give by (3, 18, 54)

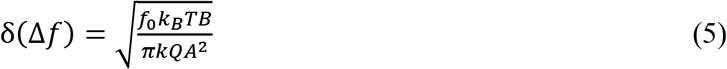

where *k_B_* is the Boltzmann constant, *T* is the absolute temperature, *B* is the measurement bandwidth (i.e., the temporal resolution), and *A* is the oscillation amplitude of the cantilever. The cantilever mass sensitivity *S_m_* is defined as (3, 18, 54)

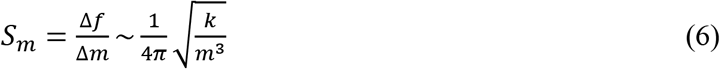

where *m* is effective mass of the cantilever. The minimum detectable mass δ(Δ*m*) can be then calculated by

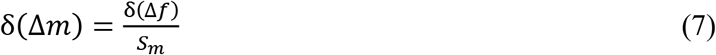

Modifying the cantilever design, *e.g.* selection of the cantilever material and dimensions, can result in substantial improvement in δ(Δ*m*). ***Fig. 6*** shows the calculated δ(Δ*m*) of cantilevers made from different materials normalized to the δ(Δ*m*) of the Si cantilever, i.e. δ(Δ*m*)/δ(Δ*m*)Si, based on **Eq. 5-7**, predicting the effect of using various candidate materials for the cantilever while keeping the dimensions and medium unchanged. The candidate materials considered included semiconductor materials like Si and Si3N4, and polymeric materials like polyimide and SU8. Parameters such as *f_o_*, *Q* and *A* for the calculation of δ(Δ*m*) were extracted from COMSOL simulation results using the model shown in ***Fig. 5(a)***. The finite element analysis was performed assuming conventional solid cantilevers. The oscillation amplitude *A* was based on a generic driving force that was kept the same for all materials simulated. Therefore, the normalized quantity δ(Δ*m*)/δ(Δ*m*)_Si_ was used for the comparison of different materials and geometries. Given the constrains mentioned above, the results obtained provide insights and design suggestions for microfluidic cantilevers operating in liquid for mass sensing purposes.

As shown in ***Fig. 6***, δ(Δ*m*) highly depends on the materials selected, and also the properties of surrounding medium with values in water significantly increased (degraded) from those in air. Polymers like polyimide and SU8 appear to provide smaller (better) detectable mass when placed in water. These polymer materials showed a 3× improvement in terms of minimum detectable mass in water compared to Si and 4.5× compared to Si_3_N_4_ (***Supplementary Information 8***). This may be related to the smaller *k* and thus flexibility of the device that leads to larger vibration amplitude *A* in water. However, polymers are not without drawbacks. Their volume could vary due to swelling in liquid(55, 56); also stiffening and softening of the polymer cantilever might result from the applied pressure in the cavity channel, as well as from potential temperature variations over long measurements. More research is required to address swelling in liquids with possible solution the deposition of very thin insulating layers by atomic layer deposition.

**Fig. 6.**
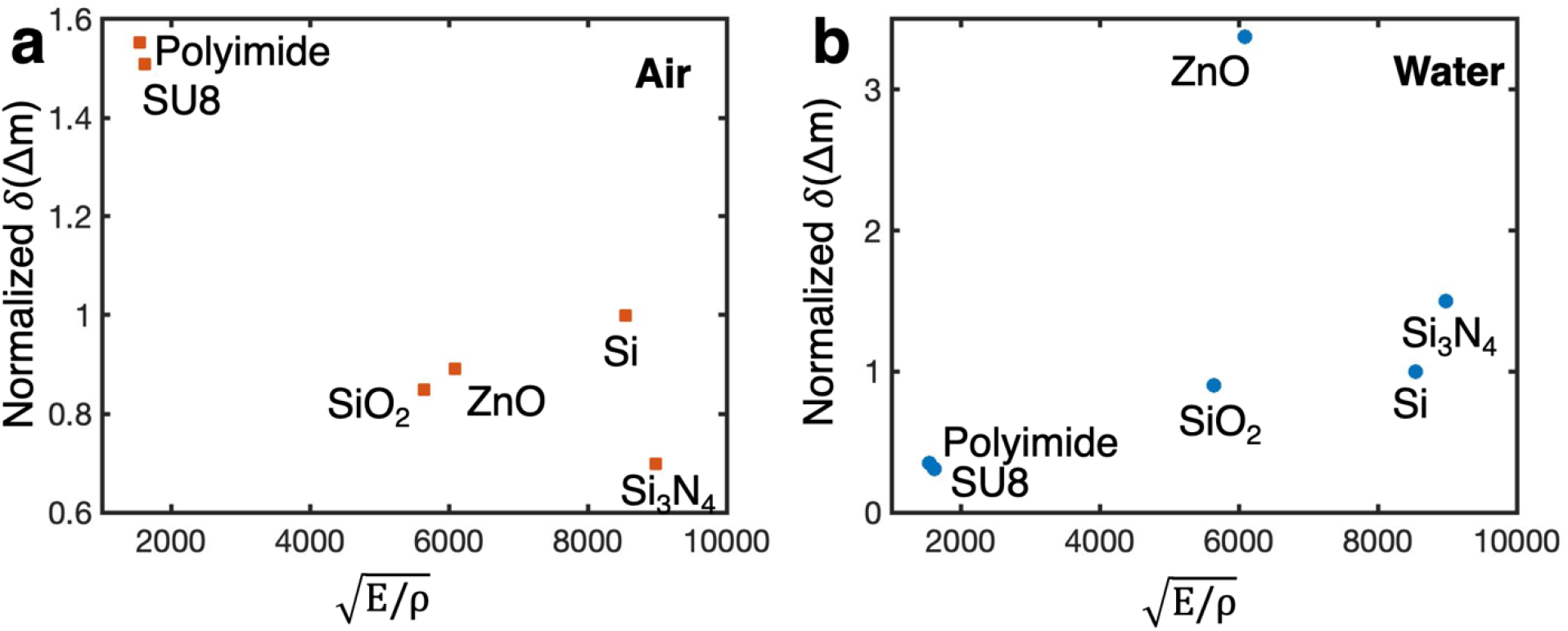
Normalized minimum detectable mass δ(Δ*m*)/δ(Δ*m*)_Si_ for cantilevers of the same size but made from different materials while resonating a) in air, and b) in water. Minimum detectable mass of Si cantilever δ(Δ*m*)_Si_ is used for normalization.

The effects of dimensions (length, width and thickness) on the minimum detectable mass δ(Δ*m*) are illustrated in ***Fig. 7***, and the effects of dimensions on the minimum detectable frequency shift δ(Δ*f*) and the mass sensitivity S_m_ are summarized in ***Supplementary Information 9***. The measurement bandwidth B in **Eq. 5** was assumed to be 100Hz based on a 10 ms time resolution. The results indicate that δ(Δ*m*) increases (degrades) with larger length, width, and thickness, while the width shows smaller effect compared to length and thickness. The same results should be applicable to different driving force. The trends are useful to help select the dimensions and materials of future cantilever designs. In general, to detect smaller mass it is desirable to use cantilevers with smaller length, smaller width, and thinner thickness.

**Fig 7:**
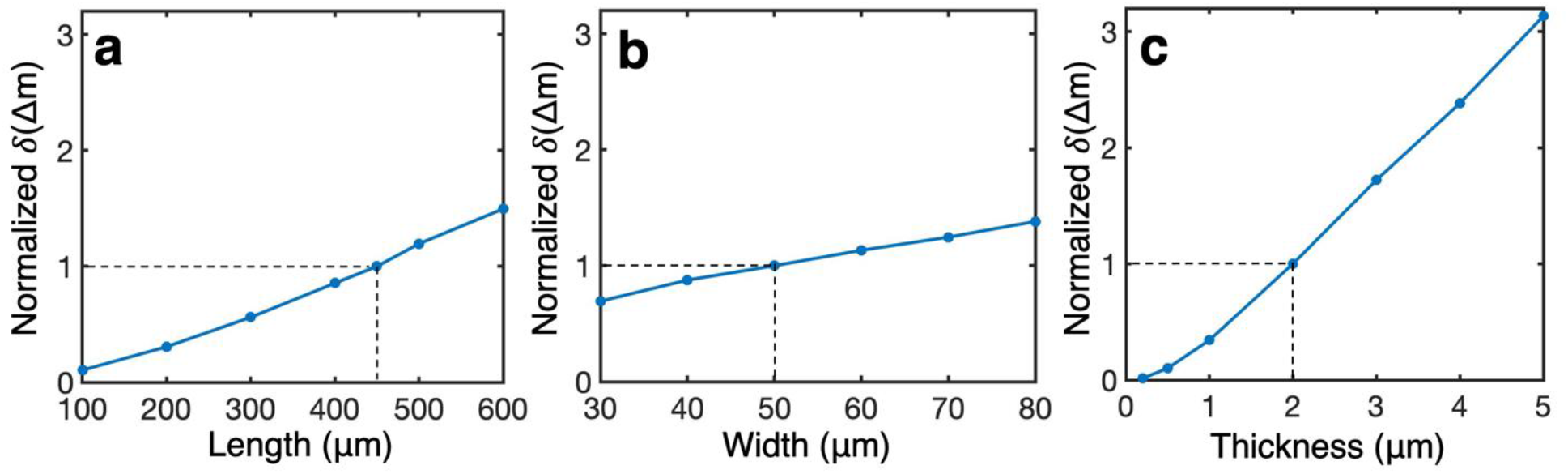
Normalized minimum detectable mass δ(Δ*m*)/δ(Δ*m*)_ref_ in water for cantilevers of different dimensions. δ(Δ*m*)_ref_ is the minimum detectable mass for a 450 μm × 50 μm × 2 μm silicon cantilever in water. While varying a) length, b) width, and c) thickness, other dimensions were kept unchanged at the nominal values of the reference cantilever. The dotted lines show the reference dimensions.

## 3. Conclusions

This work demonstrates that microfluidic cantilevers can be used for fast cellular mass measurements. We have shown the ability to measure small masses and to track changes in real-time. Despite the measurement being performed in liquid with reduced Q factor, by using the thermal noise method, instead of active vibration, and using an approximation method (22) to calculate mass from the frequency change, accurate measurements down to one hundred picograms of cell mass in a physiologically relevant environment were obtained. Finite element analysis suggests that polymer materials would allow for a significant improvement in minimum detectable mass compared to the conventional microcantilever materials used in our studies. Tailoring microcantilever dimensions by having smaller length, width, and thickness would also improve performance. Instead of the thermal tune method (49, 50); we can actively vibrate the cantilever to increase the amplitude of oscillation and thus decrease the minimum detectable mass. Exciting the piezoelectric scanners of the AFM to drive the cantilever in liquids creates forest of peaks (57–59), which make it hard to distinguish the resonance frequency. There are several techniques that can be used to address this issue, such as using a shock absorbing material (59) and bringing the piezoelectric actuator very close to a conventional cantilever (60). Another method is to thermally induce stress with a laser near the fixed end of the cantilever (photo-thermal modulation)(61, 62). Other methods include cantilevers with integrated piezoelectric actuation (63–65), cantilevers with integrated thermomechanical actuation (66–68), using magnetic excitation (69–71), or a measurement strategy based on phase-locked loop (72).

## Acknowledgments

We would like to thank Prof. Stuart Sealfon (SCS) for fruitful discussions and advice, Dr. Edward Nelson for advice with AFM, Mr. Sagie Mofsowitz for help with cell culture, for Ni-ka Ford help with *Fig. 1a* graphics. This work was supported in part by the National Institutes of Health (U24DK112331-03S1 Diversity Supplement (SCS, AG), R01AI121012 (BG), R21AI137785 (BG and AG), R44GM146477 (AG)). The opinions expressed here are those of the authors and do not represent the official position of the NIH or the United States Government.

## Data access statement

All data that support the findings of this study are included within the article (and any supplementary information files).

## Ethics statement

This work does not include human subjects, human data, or animals.

