## Supplemental material for "AFM Microfluidic Cantilevers as Weight Sensors for Live Single Cell Mass Measurements"

**1. Uncertainties in defining the spring constant**

| | Experiment 1<br>Spring Constant (N/m) | Experiment 2<br>Spring Constant (N/m) |
|---|---|---|
| Run 1 | 2.139 | 2.077 |
| Run 2 | 2.237 | 2.214 |
| Run 3 | 2.165 | 2.043 |
| Run 4 | 2.123 | 2.031 |
| Average | 2.166 | 2.091 |
| STD | 0.05 | 0.08 |

Table SI1: The spring constant measured of a tipless fluidic cantilever, Probe 16730 (Cytosurge, AG, Switzerland). Manufacturer provided its spring constant being 2N/m, with a 4 micron aperture. The fluidic cantilever is filled with DI water, and the spring constant is measured in air with thermal tuning method. Differences of the spring constant obtained across experiments can be observed, which could result in an error in the subsequent mass measurements.

### 2. Uncertainties in measuring the natural frequency in liquid

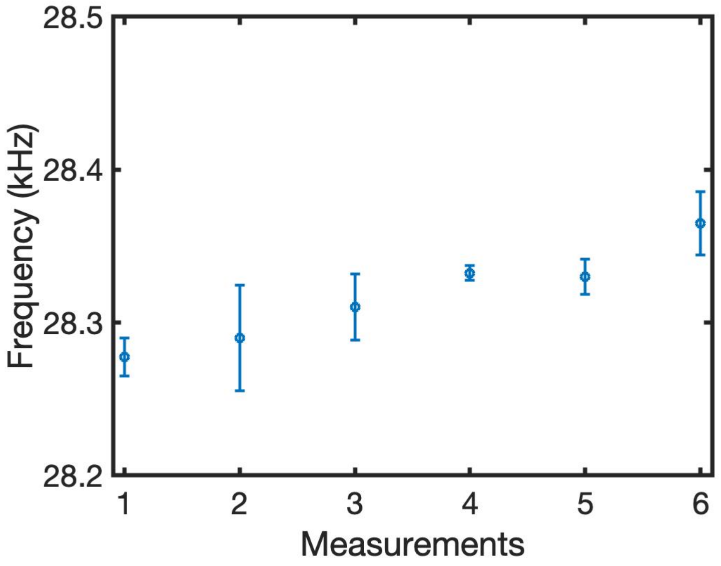


Figure SI2: We observed that changes could happen in the resonance frequency $f_0$ while conducting experiments in liquid.  Figure SI3 shows an example of resonance frequencies fluctuations range from 28.28 kHz to 28.37 kHz, or a shift of 0.9 kHz within six experiments. Every measurement of resonance frequency is ran 4 times to get an average, and each run the measurements are averaged over 250 iterations.

### 3. Information on growth media

The growth media compositions are 500mL Dulbecco's Modified Eagle's Media (DMEM) (Corning, USA) supplemented with 4.5 g/L glucose, L-glutamine, and sodium pyruvate, 50mL Fetal Bovine Serum (FBS) (Thermal Fisher, USA), and 5mL Penicillin-Streptomycin (Thermal Fisher, USA) The HEK293 cells are kept in this growth media at 37°C in a humidified atmosphere with 5% $CO_2$. We use 0.25% trypsin-EDTA (Thermal Fisher, USA) to detach the HEK293 from the culture dish and add in the 1mL of HEPES buffer (Corning, USA) into 50mL of growth media to create a 20mM HEPES growth media solution for subsequent experiments with the AFM.

**4. No distinguishable effect of applied pressure on the measured frequency of the cantilevers**

| | **Frequency at 20mbar (kHz)** | **Frequency at -300mbar (kHz)** |
|---|---|---|
| Run1 | 28.43 | 28.41 |
| Run2 | 28.39 | 28.4 |
| Run3 | 28.41 | 28.45 |
| Run4 | 28.44 | 28.41 |
| Average | 28.42±0.02 | 28.42±0.02 |

Table SI4: An example of the measured resonance frequency $f_0$ in liquid at 20mbar and -300mbar. At 20mbar, the cantilever's resonance frequency is 28.42±0.02 kHz, and when switched to -300mbar, the measured resonance frequency is 28.42±0.02 kHz. There are no distinguishable differences at the measured frequency when the pressure applied is varied.

5. **Cell tracker green dye for HEK293 cell at -100mbar pressure**

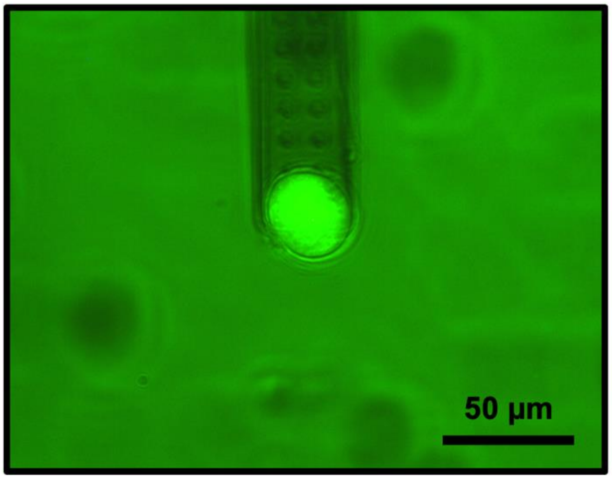


FigureSI5. A fluorescence image of HEK293 cell using Celltracker Green (Invitrogen$^{TM}$) attached by applying 100 mbar of negative pressure on the fluidic cantilever.

When leaving the cell on the cantilever with -100mbar of negative pressure, we do not observed the fluorecent dyes migrating into the cantilever. Thus we ruled out the possibility of the negative pressure breaking the cellular membrane and drawing the cytoplasm from the cell.

**6. Evaporation of the media for long experiments**

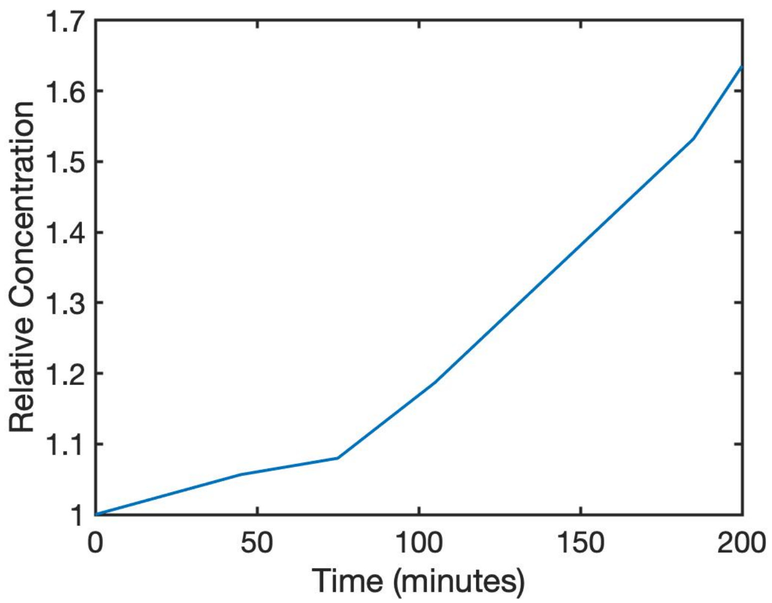


FigureSI6: Relative concentration of media measured across time. At the starting time of 0 minutes, we normalize the concentration to 1. Due to the constraints of the incubator set-up without a moisture control, we observe that the evaporation results in the increase of media concentration is about 10% in the first hour of measurement, yet the loss of media could be adjusted back by adding DI water to maintain osmolarity.

### 7. Parameters for Finite element analysis

| *Parameter* | *Value* | |
|---|---|---|
| | *Air* | *Water* |
| $d_{visc}$ | 0.22[mm]*sqrt(100/$f_0$) (Hz) | 0.055[mm]*sqrt(100/$f_0$) (Hz) |
| $\mu_b$ | $1.088\times10^{-5}$ (Pa·s) | $2.81\times10^{-3}$ (Pa·s) |
| $\mu$ | $1.814\times10^{-5}$ (Pa·s) | $1.009\times10^{-3}$ (Pa·s) |
| $c$ | 343.2 (m/s) | 1481.4 (m/s) |
| $\rho$ | 1.204 (kg/m$^3$) | 998.2 (kg/m$^3$) |
| $\rho_{Si}$ | 2329 (kg/m$^3$) | |
| $E_{Si}$ | 170 (GPa) | |
| $v$ | 0.28 | |

Table SI7: Parameters used in FEA modeling with thermoviscous acoustics (frequency domain) module of COMSOL Multiphysics.

Note: $d_{visc}$ is the viscous boundary layer thickness at $f_0$; $\rho_{Si}$, $E_{Si}$ and $v$ are the density, Young's modulus, and Poisson's ratio of silicon, respectively; ρ, $\mu_b$, $\mu$, and $c$ are the density, bulk viscosity, dynamic viscosity, speed of sound of the medium, respectively.

### 8. Minimum detectable mass for cantilevers of different materials in air and water

| **Materials** | **Normalized minimum detectable mass in air** $\delta(\Delta m)/\delta(\Delta m)_{Si}$ | **Normalized minimum detectable mass in water** $\delta(\Delta m)/\delta(\Delta m)_{Si}$ | **Improvement ratio compared to Si cantilever** **(**Air / water) | **Improvement ratio compared to $Si_3N_4$ cantilver** **(**Air / water) |
|---|---|---|---|---|
| Si | 1 | 1 | 1 | - |
| $Si_3N_4$ | 0.976 | 1.5 | - | 1 |
| SU8 | 2.108 | 0.31 | 0.48 / 3.23 | 0.463 / 4.84 |
| Polyimide | 2.168 | 0.347 | 0.46 / 2.88 | 0.451 / 4.32 |

Table SI8: The minimum detectable mass for Si, $Si_3N_4$, SU8, and Polyimide in air and in water. We could see that despite the SU8 and Polyimide perform worse in air compared to Si cantilever, they can achieve a much smaller minimum detectable mass in water compared to Si, with a 3.23 and 2.88 fold improvement respectively.

### 9. Optimization of the geometry of the cantilevers

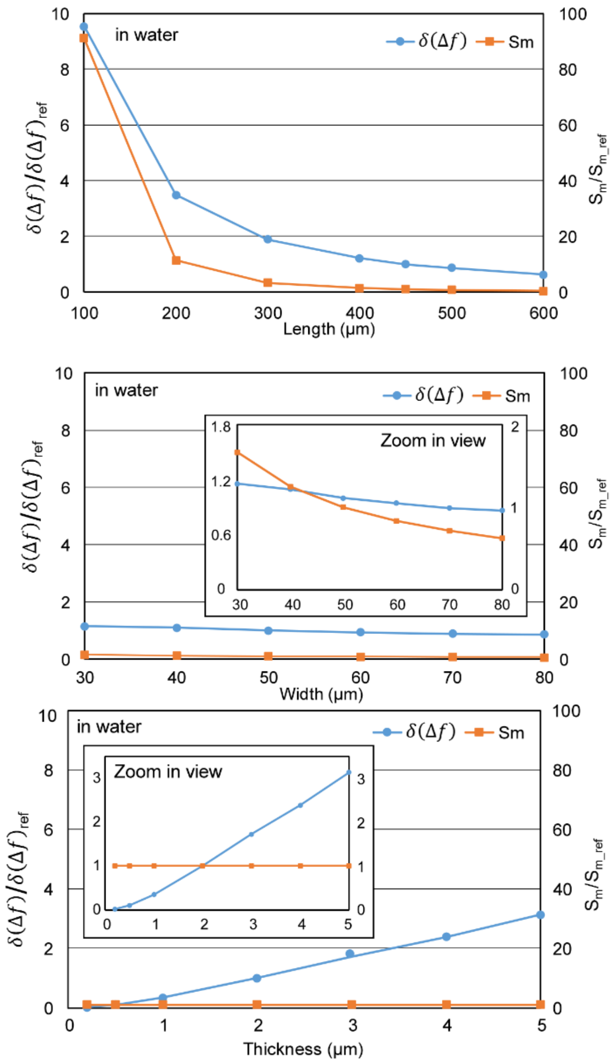


**Figure SI9.** The effects of dimensions (length, width and thickness) on minimum detectable frequency shift ($\delta(\Delta f)$) and mass sensitivity ($S_m$) of Si cantilever in water. The plots are normalized to $\delta(\Delta f)_{ref}$ and $S_{m\_ref}$ which are the minimum detectable mass and mass sensitivity, respectively, for a 450 μm × 50 μm × 2 μm silicon cantilever in water. The same y axis scale is used for a straightforward comparison between length, width and thickness. The zoom in view figures are inserted to show the details for width and thickness. The results indicate that $\delta(\Delta f)$ increases with smaller length, width and larger thickness, while $S_m$ typically increases with smaller length and width. Thickness does not affect $S_m$ as $t$ is canceled out in the calculation. Again, trade-offs between $\delta(\Delta f)$ and $S_m$ must be made. To achieve higher mass sensitivity, it is desirable to use cantilevers with higher spring constant and smaller mass (i.e., smaller length and width). To have smaller (better) frequency resolution, larger length and width are desirable as well as thinner thickness. A possible approach is to first select desired $S_m$ by determining the length and width of the cantilever, and then vary the thickness of the cantilever to adjust $\delta(\Delta f)$. The trends can be useful to help select the dimensions and materials of future cantilever designs for different sensing purposes.
